# *Mycobacterium tuberculosis* stimulates IL-1β production by macrophages in an ESAT-6 dependent manner with the involvement of serum amyloid A3

**DOI:** 10.1101/2020.04.04.025411

**Authors:** Bock-Gie Jung, Ramakrishna Vankayalapati, Buka Samten

**Author notes:** Corresponding Author mailing address: Department of Pulmonary Immunology, The University of Texas Health Science Center at Tyler, 11937 US Hwy 271 Tyler, Texas 75708-3154.

## Abstract

To explore interleukin (IL)-1β production in tuberculosis, we infected mouse bone marrow-derived macrophages (BMDM) with *Mycobacterium tuberculosis* (*Mtb)* H37Rv, its early secreted antigenic target protein of 6 kDa (ESAT-6) gene deletion (H37Rv:Δ3875) or complemented strain (H37Rv:Δ3875C) and evaluated IL-1β production. H37Rv induced significantly increased IL-1β production by BMDMs compared to non-infected BMDMs. In contrast, H37Rv:Δ3875 induced significantly less mature IL-1β production despite eliciting comparable levels of pro-IL-1β and IL-8 from BMDMs compared to H37Rv and H37Rv:Δ3875C. Blocking either NLRP3 or K^+^ efflux diminished H37Rv-induced IL-1β production by BMDMs. Infection of mice intranasally with H37Rv:Δ3875 induced less IL-1β production in the lungs compared with H37Rv.Intranasal delivery of ESAT-6 but not CFP10 induced production of IL-1β in mouse lungs and RNA-Seq analysis identified serum amyloid A (SAA) 3 as one of the highly expressed genes in mouse lungs. Infection of mice with H37Rv but not H37Rv:Δ3875 induced expression of lung SAA3 mRNA and protein, consistent with the effect of intranasal delivery of ESAT-6. Silencing SAA3 reduced *Mtb-*induced IL-1β production by BMDMs. We conclude that the production of SAA3 is required for *Mtb* stimulated IL-1β production by macrophages in tuberculosis infection.

## INTRODUCTION

Tuberculosis (TB) caused around 1.2 million deaths among HIV-negative people and an additional 251,000 deaths among HIV-positive people in 2018 (1), which is mainly due to the lack of a detailed understanding of the molecular mechanisms of interactions between immune cells and the pathogen, *Mycobacterium tuberculosis* (*Mtb*). Improved understanding of the host-pathogen interaction will lead to an effective TB vaccine design and an identification of novel drug targets to improve clinical management of drug-resistant TB infections. Macrophages play critical roles in tuberculosis infection as both host cells providing intracellular niche for *Mtb* infection and growth and effector immune cells fighting against *Mtb* infection by communicating with other immune effector cells by producing different cytokines (2).

Although IL-1β production by macrophages is pivotal for protection against TB infection (3, 4), several clinical reports showed that IL-1β is also involved in TB pathogenesis. There is significantly increased IL-1β mRNA in the alveolar macrophages and IL-1β protein in bronchoalveolar lavage fluid of TB patients compared with healthy controls (5). Patients with TB pleurisy also had higher IL-1β in their serum and pleural fluid than patients with lung cancer (6). Moreover, high-IL-1β expressing genotype correlated with disease progression and poor treatment outcomes of TB patients (7). These observations suggest that IL-1β correlates with the severity of TB. Therefore, it is critical to delineate mechanisms of *Mtb* induced IL-1β production and its role in TB pathogenesis.

IL-1β production by immune cells requires two signals, priming and activation. The priming initiated by pattern recognition receptors (PRRs) upon recognition of pathogen-associated molecular patterns (PAMPs) leads to transcription of IL-1β and related genes in NF-κB (8) and STAT3 (9) dependent pathways and production of pro-IL-1β and inflammasome components; the activation provided by intracellular PRRs and damage-associated molecular patterns (DAMPs) initiates the assembly of inflammasome and activation of caspase-1 which cleaves pro-IL-1β into mature IL-1β (8) and cleaves gasdermin D to form cellular membrane pores for the release of mature IL-1β (10). Inflammasomes are activated by either PAMPs or DAMPs directly (11) or indirectly through P2×7 receptor (12). Serum amyloid A protein (SAA)3, the most predominant isoform in the lung (13), activates nod-like receptor protein (NLRP)3 inflammasome in a cathepsin B-sensitive pathway *via* P2×7 receptor (14) and induces mature IL-1β production by macrophages. Potassium (K+) efflux is one of the essential downstream triggers for inflammasome activation and mature IL-1β production (15, 16).

*Mtb* infection of macrophages induces the production of mature IL-1β through the early secreted antigenic target of 6-kDa (ESAT-6) dependent activation of NLRP3 with the involvement of cathepsin B (17, 18). However, the role of SAA3 in IL-1β production by macrophages in tuberculosis infection remains unexplored. Using both in vitro and in vivo studies, we demonstrated in this study that *Mtb* stimulates mature IL-1β secretion by macrophages *via* ESAT-6 dependent expression of SAA3 and activation of NLRP3 and K^+^ efflux.

## MATERIALS AND METHODS

### Animals

Female 6-8-week old C57BL/6 mice were from The Jackson Laboratory. All the animal studies were performed in compliance with the NIH guidelines and regulations and approved by the Institutional Animal Care and Use Committee of the University of Texas Health Science Center at Tyler.

### Mouse bone marrow-derived macrophages

Mouse bone marrow-derived macrophages (BMDMs) were prepared as previously described with minor modifications (19).

### *Mtb* strains

*Mtb* strain H37Rv, its *esat*-6 deletion mutant (H37Rv:Δ3875), or *esat*-6 complemented strain (H37Rv: Δ3875C) were cultured and prepared for infection, as previously described (20).

### Recombinant m*ycobacterial proteins*

Recombinant ESAT-6 and culture filtrate protein of 10 kD (CFP10) were expressed in *E. coli* and prepared as described earlier (19, 21), aliquoted, and stored at - 80 °C before use.

### *Intranasal Mtb* infection or recombinant proteins inoculation of mice

The mice with complete anesthetization were intranasally administrated 50 µl of live *Mtb* and determined to be 30,000 colony forming units (cfu) of bacilli per mouse as determined one day after infection. The recombinant proteins were administrated intranasally, as previously described (19).

### Lung *Mtb* burden

Bacilli burden in the lungs at 3, 7, 10, and 14 days post-infection (DPI) was determined by culturing serially diluted mouse lung homogenates on 7H10 agar plates (BD Biosciences) at 37°C for 2-3 weeks. The colonies were counted and expressed as log cfu/ml.

### ELISA

Levels of IL-1β, KC and IL-6 in the cell culture supernatants and lung homogenates were determined by commercialized ELISA kit (IL-1β and KC were from R&D Systems BD, IL-6 was from Biolegend) and SAA3 in the lung homogenates and plasma were determined by ELISA kit (MilliporeSigma) following the manufacturer’s instructions.

### Western blot

Levels of pro-IL-1β, IL-1β and KC in the cell lysates and the culture supernatants were determined by western blot analysis using glyceraldehyde 3-phosphate dehydrogenase (GAPDH) as loading control with specific antibodies (Cell Signaling Technology) as previously described (19).

### RNA-Sequencing

At one day after intranasal inoculation of ESAT-6 or culture filtrate protein 10 kDa (CFP10), the total RNA was extracted from the mice lungs using Trizol LS reagent (Invitrogen), and the samples passed RNA quality tests were used for RNA-Seq analysis. The paired-end mRNA sequencing was performed on the Illumina platform as per the standard protocols at LC Science (Houston, USA). The reads were mapped to mouse genome using Bowtie 2. Transcript abundances were estimated using RSEM v1.3.0. Differential expression analysis was performed using EdgeR in the Bioconductor. Genes showing significant differences (FDR < 0.05) were selected for enrichment analysis using GAGE v2.20.1.

### Real-time PCR

The expression of mRNA levels of IL-1β and SAA3 in BMDMs and lung homogenate cells was measured by Real-Time PCR from the total RNA using GAPDH as internal control with specific primer and probe sets (Applied Biosystems) as previously described (19).

### Transfection of siRNA

BMDMs with 80% confluence were transfected with either scrambled or two different concentrations of SAA3 siRNA (Cell Signaling Technology) using Lipofectamine RNAiMAX reagent (Life Technologies) following the manufacturer’s instructions. The cells transfected with siRNA for 24 h were infected with *Mtb* H37Rv at 10 multiplicity of infection (MOI) for 24 h. The levels of IL-1β and IL-6 in the culture supernatants were determined by ELISA. The silencing efficiency was determined by evaluating SAA3 mRNA levels in siRNA transfected cells by real-time PCR.

### Statistical analysis

Data are expressed as mean ± standard error of the mean. Student’s t-test was performed for statistical analysis using GraphPad Prism version 4.0 software (GraphPad Software, Inc.). A P value of less than 0.05 was considered statistically significant.

## RESULTS

### *Mtb* induces mature IL-1β production by macrophages in an ESAT-6 dependent manner

To test the role of ESAT-6 in *Mtb* induced IL-β production, mouse BMDMs were infected with H37Rv, H37Rv:Δ3875 or H37Rv:Δ3875C followed by determination of pro- and mature IL-1β and keratinocyte chemoattractant (KC) in the culture supernatants by ELISA and western blotting, respectively. H37Rv induced significantly increased levels of IL-1β compared to uninfected BMDMs, and the majority of the IL-1β in the culture supernatant was 17 kDa mature IL-1β as determined by western blot analysis. In contrast, H37Rv:Δ3875 induced significantly less IL-1β production (Fig. 1A) and complementation of *esat-6* rescued mature IL-1β production (Figs. 1A and C). All three strains of *Mtb* induced comparable levels of both KC (Fig. 1B) and pro-IL-1β (Fig. 1D) and further incubation of the infected BMDMs with ATP, an NLRP3 activator, induced comparable levels of mature IL-β production by all three strains (data not shown), suggesting that the reduced IL-1β production by H37Rv:Δ3875 is not due to less priming of pro-IL-1β because of reduced infectivity. These data indicated that ESAT-6 is essential for macrophage production of mature IL-1β in response to *Mtb* infection, consistent with previous reports (17, 22).

**Figure 1.**
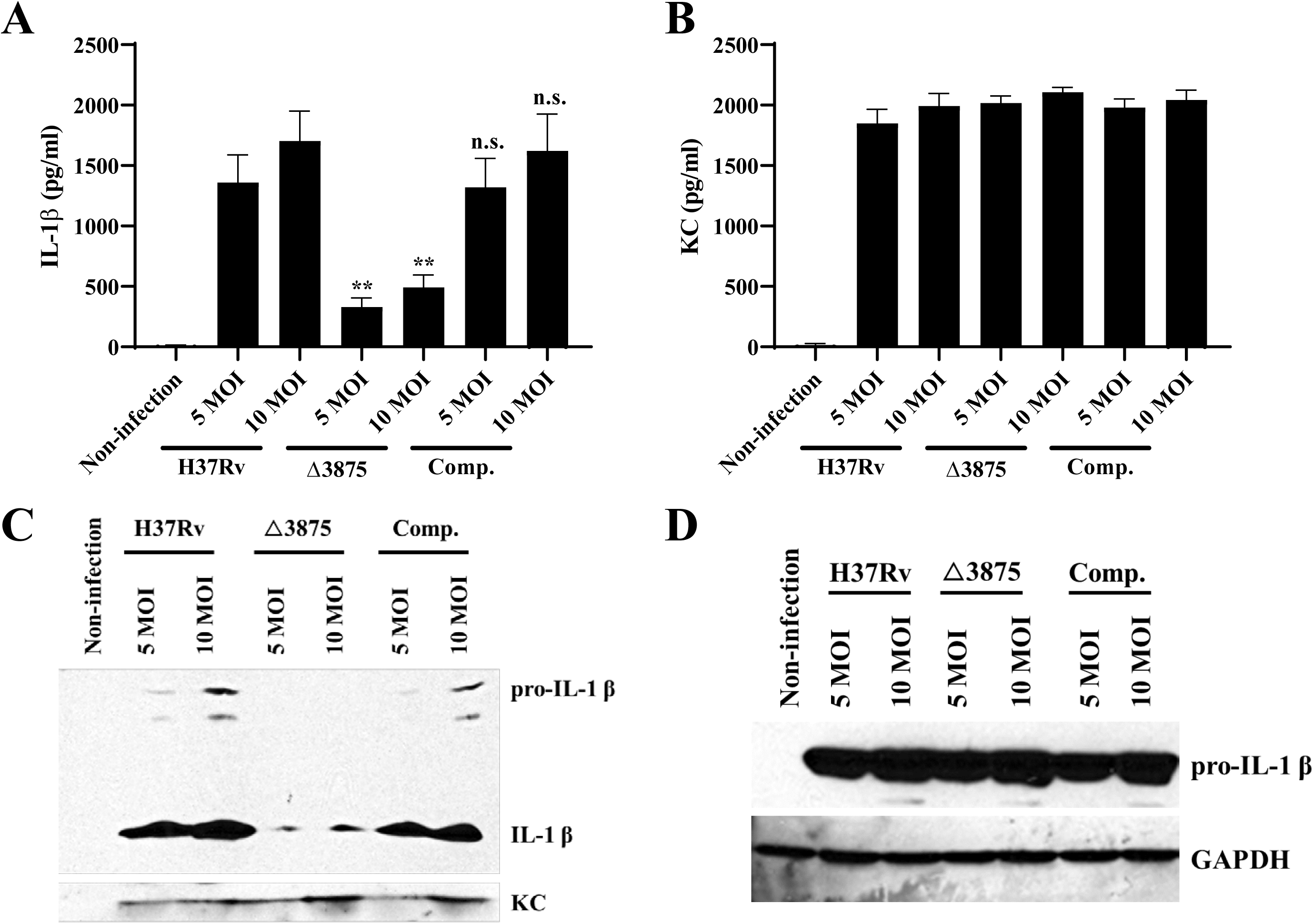
*Mtb* infection induces maturation and secretion of IL-1β by macrophages in an ESAT-6 dependent manner. Mouse bone marrow-derived macrophages (BMDMs) were infected with *Mtb* strain H37Rv, its *esat*-6 deletion mutant (Δ3875), or *esat*-6 complemented strain (Comp.) at 5 and 10 MOI. At 24 h post-infection, the levels of IL-1β (A) and KC (B) were determined by ELISA and the levels of pro-IL1β, mature IL-1β and KC in the cell culture supernatants were determined by western blot (C). At 12 h post-infection, the levels of pro-IL-1β in the cell lysates were determined by western blot and the membrane was stripped and blotted for GAPDH as loading control (D). Data are expressed as Means ± SEM. n.s., not significant; *, P < 0.05; **, P < 0.01. Western blot data are one representative result of 3 independent experiments.

### *Mtb* induces mature IL-1β production by macrophages through NLRP3 inflammasome

Since NLRP3 is the major inflammasome for caspase-1-mediated IL-1β maturation in *Mtb*-infected macrophages (17), we tested the role of NLRP3 in our system. Pretreatment of BMDMs with MCC950, an NLRP3 specific inhibitor (23, 24), suppressed mature IL-1β production by H37Rv infected BMDMs dose-dependently (Fig. 2A) without affecting the expression of cellular pro-IL-1β by BMDMs (Fig. 2B), confirming the critical role of NLRP3 inflammasome in *Mtb* induced mature IL-1β production by macrophages. Cellular K^+^ efflux through TWIK2 potassium channel (25) activates inflammasome for the maturation of IL-1β in macrophages infected with intracellular bacteria (26, 27) by acting as a downstream inflammasome activator (28). Thus, we also tested the significance of this mechanism in *Mtb*-induced IL-1β production by incubating H37Rv infected BMDMs with different concentrations of KCl, quinine, a specific inhibitor of TWIK2 K^+^ efflux channel (25) or NaCl as control. Consistent with previous studies (25–27), IL-1β secretion was inhibited by increased extracellular KCl and quinine, but not by NaCl (Fig 2C), implying that K^+^ efflux through TWIK2 K^+^ efflux channel is required for *Mtb*-induced IL-1β production. This was not due to either reduction of pro-IL-1β (data not shown) or general cytotoxicity because neither of them affected cell viability as determined by MTT assay (Fig. 2D).

**Figure 2.**
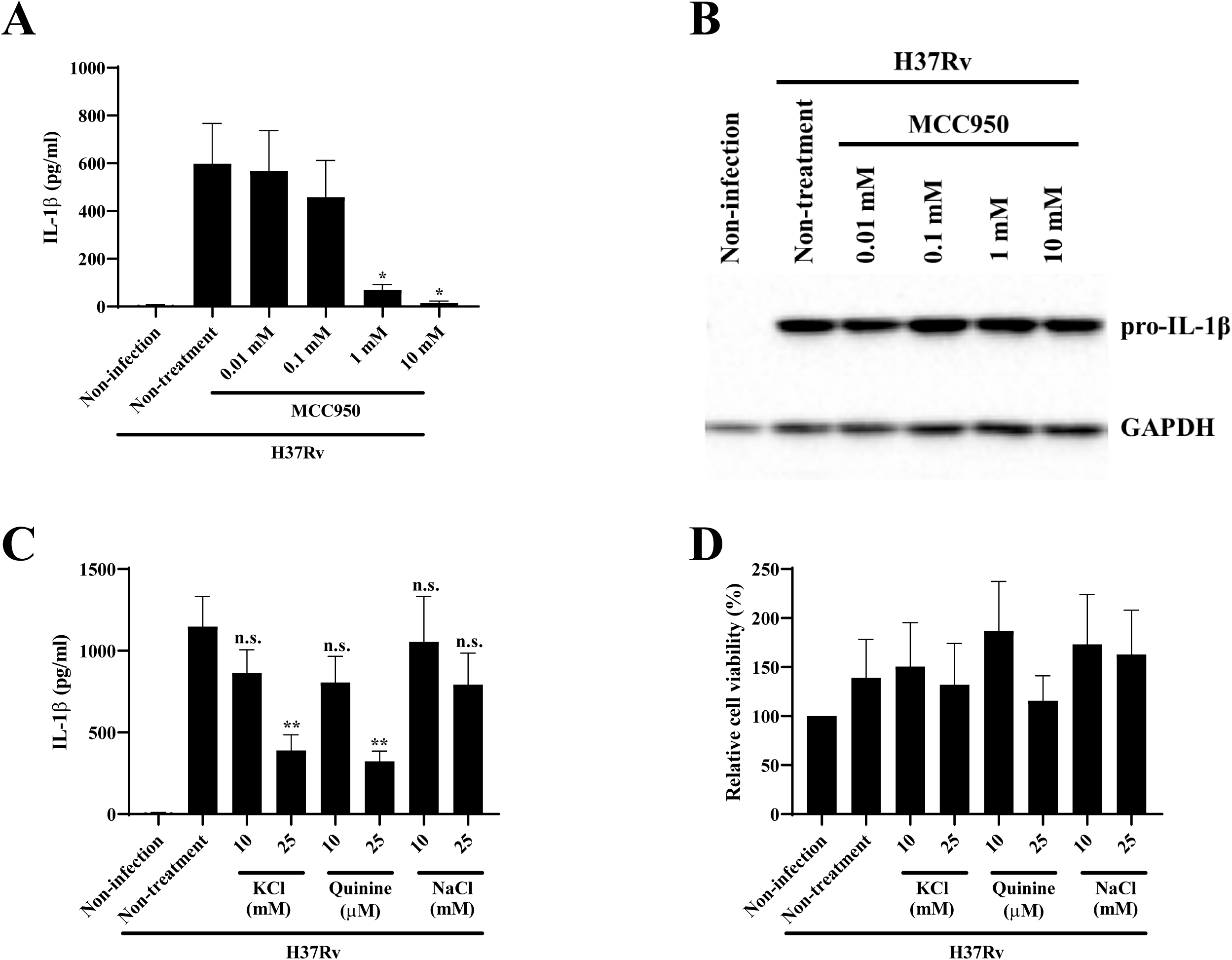
*Mtb* induces maturation and secretion of IL-1β by macrophages through NLRP3 inflammasome activation and potassium efflux mechanisms. BMDMs were pretreated with different concentrations of MCC950, an NLRP3 inhibitor, and infected with H37Rv for 24 h. IL-1β (A) in the culture supernatants was determined by ELISA and pro-IL-1β (B) in the cellular protein lysates were determined by western blot, respectively. BMDMs were infected with H37Rv at 10 MOI for 24 h in the presence of different concentrations of KCl, quinine or NaCl. The levels of IL-1β (C) in the cell culture supernatants were determined by ELISA. Cell viability (D) was evaluated by MTT assay. Data are expressed as Means ± SEM. n.s., not significant; *, P < 0.05; **, P < 0.01. Western blot data are one representative result of 3 different experiments.

### ESAT-6 is required for *Mtb* induced IL-1β production

To confirm our *in vitro* data in an *in vivo* infection model, we infected mice with a high dose *Mtb* (H37Rv) as H37Rv:Δ3875 shows growth defects in mice (29). Interestingly, despite the delivery of 30,000 cfu of *Mtb* into the mouse lungs and this led to 40-50 thousand bacilli of both strains in mouse lungs at 3 and 7 DPI (Fig. 3B), IL-1β levels in the mouse lungs remained similar to that of mock infection (HBSS control, data not shown). A significant increase of lung IL-1β was evident only at 10 DPI in H37Rv infected mice and reached significantly higher levels at 14 DPI compared with that of H37Rv:Δ3875 infected mice (Fig. 3A). Though both strains grew at a similar rate from 1-7 DPI, the growth of H37Rv increased significantly higher than that of H37Rv:Δ3875 at 14 DPI (Fig. 3B). These data imply that despite higher inoculation dose, *Mtb* does not stimulate significant amount of lung IL-1β production and inflammation until 10-14 DPI, consistent with the inflammatory responses of the mice infected with low dose aerosol *Mtb* (30), suggesting that the bacilli number at least up to 30,000 CFU is not the determining factor for the initiation of IL-1β production and immune responses in mouse lungs and ESAT-6 is required for *Mtb* induced IL-1β production *in vivo*. To address whether the reduced production of IL-1β in mouse lungs by H37Rv:Δ3875 is due to lack of ESAT-6, but not simply due to reduced bacilli number, we intranasally delivered ESAT-6, CFP10 or HBSS (vehicle control) and determined the lung IL-1β mRNA and proteins at one-day post-treatment. ESAT-6 but not CFP10 nor HBSS induced significantly increased IL-1β mRNA (Fig. 3C) and protein (Fig. 3D), indicating that ESAT-6 but not CFP10 induces IL-1β production *in vivo*. This is less likely due to LPS contamination of ESAT-6. Because incubation of BMDMs with ESAT-6 or LPS resulted in a different activation pattern of signaling molecules in macrophages, including phosphorylation of mitogen activated protein kinases (MAPK) and signal transducer and activator of transcription (STAT) 1 and 3. Although both stimulants induced activation of all three members of MAPK, they differ strikingly in activations of STAT1 and STAT3. ESAT-6 but not LPS induced activation of both STAT1 and STAT3 (Fig. 3E) consistent with our published work (19) and consistent with previous study that LPS induces IL-1β gene expression *via* NF-κB but not STAT3 pathway (9).

**Figure 3.**
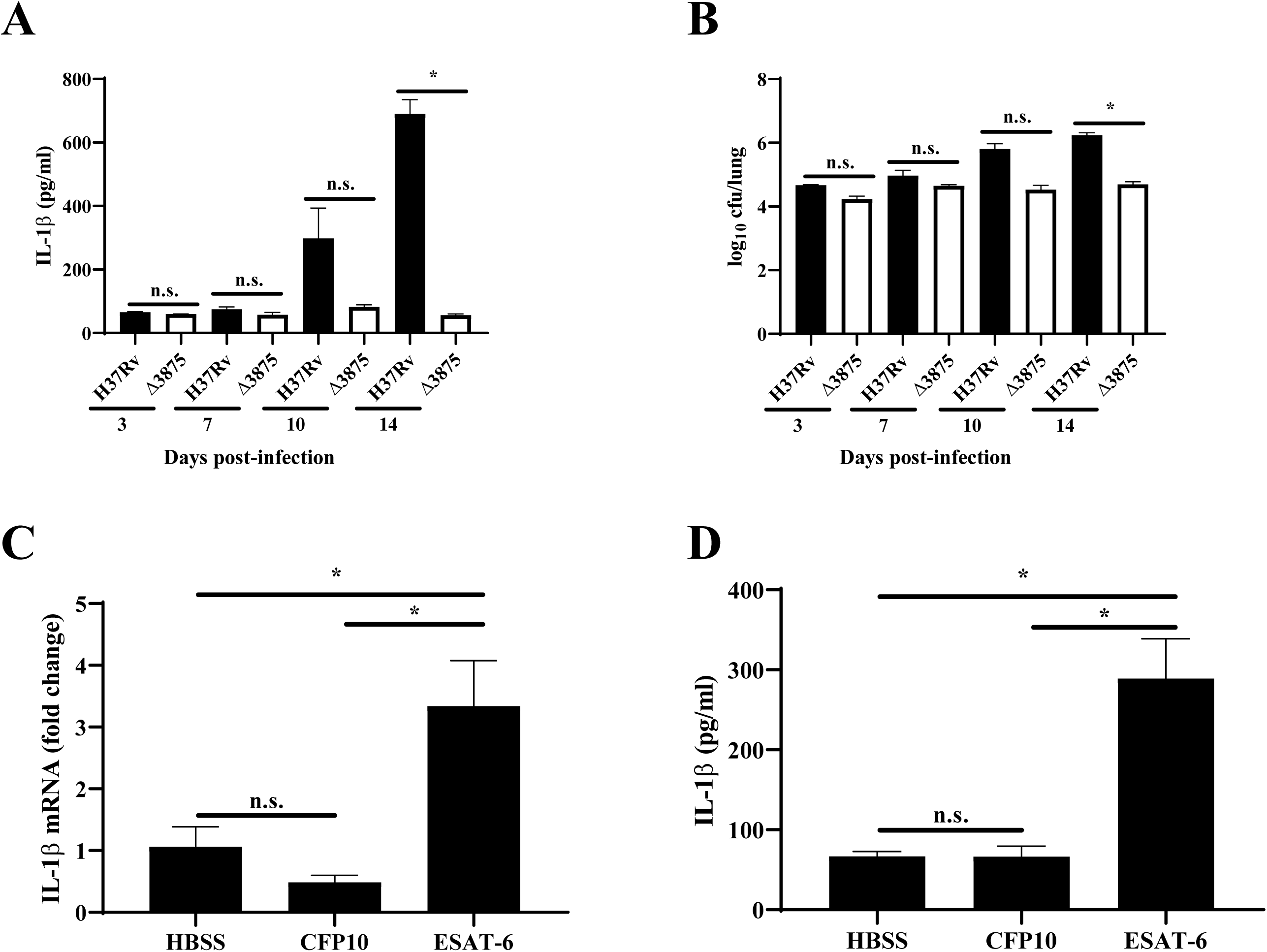
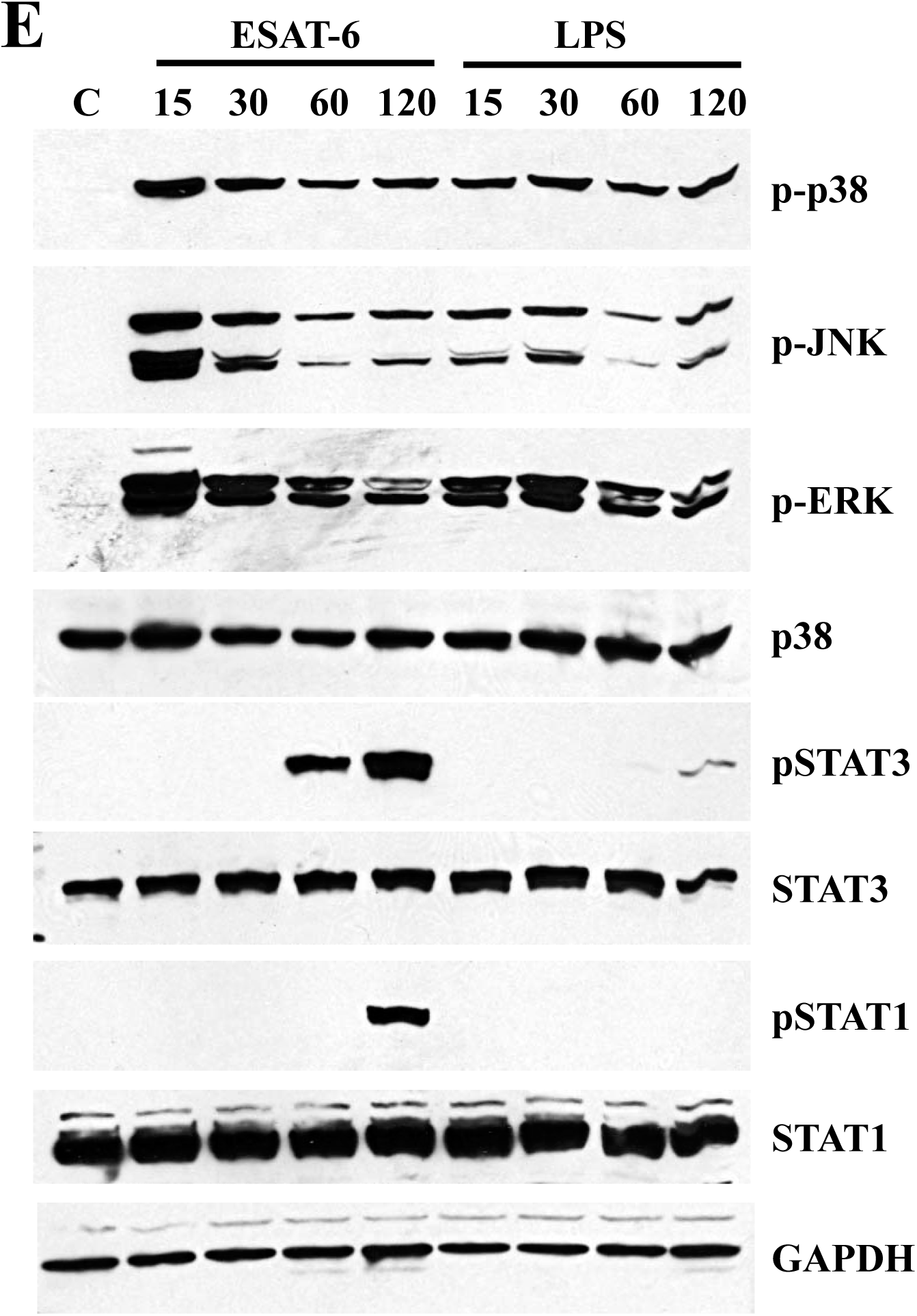
Mtb stimulates IL-1β production in mouse lungs in an ESAT-6 dependent manner. C57BL/6 mice were infected intranasally with H37Rv or its *esat*-6 deletion mutant H37Rv: Δ3875 (Δ3875) to deliver 30,000 colony-forming units (cfu) and sacrificed at 3, 7, 10, and 14 days post-infection. The levels of IL-1β (A) and cfu of *Mtb* (B) in lung homogenates were determined. C57BL/6 mice were treated intranasally with ESAT-6 or CFP10 at 40 µg/mouse or equal volume of HBSS as vehicle control) and mRNA (C) and protein levels (D) of IL-1β in the lung homogenates at one day post inoculation were determined by real-time PCR and ELISA, respectively. BMDMs were incubated with or without ESAT-6 or LPS until 120 min and phosphorylation of indicated kinases and signal transducer and activator of transcriptions (STAT) were evaluated in cell lysates by western blot (E). Data are expressed as Means ± SEM. n = 3, one of three independent experiments. n.s., not significant; *, P < 0.05. Western blot data are one representative result of 4 different experiments.

### Intranasal inoculation of ESAT-6 upregulates SAA3 expression in mouse lung

To explore the potential effects of ESAT-6 on inflammatory responses in vivo, we determined mouse lung gene-expression profile by RNA-sequencing analysis of total lung RNA of mice inoculated intranasally with ESAT-6 or CFP10. The RNA-sequencing analysis identified a total of 587 out of 19,109 mouse genes as differentially expressed between ESAT-6 and CFP10 inoculated mice. Out of these, ESAT-6 upregulated 443 genes and downregulated 144 genes compared with CFP10. Gene set pathway enrichment analysis identified that the majority of the differentially expressed genes belongs to the signaling pathways of IL-17, cytokine-cytokine receptor interaction, NOD-like receptor, TNF, Toll-like receptor, chemokine, Jak-STAT and osteoclast differentiation with gene numbers for each pathway ranging from 14-42 (Fig. 4A). Interestingly the majority of these genes were also expressed by macrophages upon infection with a hypervirulent *Mtb strain* (30, 31). The top 20 genes with the highest p values were shown with the heat map demonstrated that SAA3 is one of the most highly upregulated genes by ESAT-6 (Fig. 4B). This was further confirmed by significantly upregulated expression of SAA3 mRNA (Fig. 4C) and protein (Fig.4D) in mouse lungs inoculated with ESAT-6 as determined by real-time PCR and western blotting, respectively.

**Figure 4.**
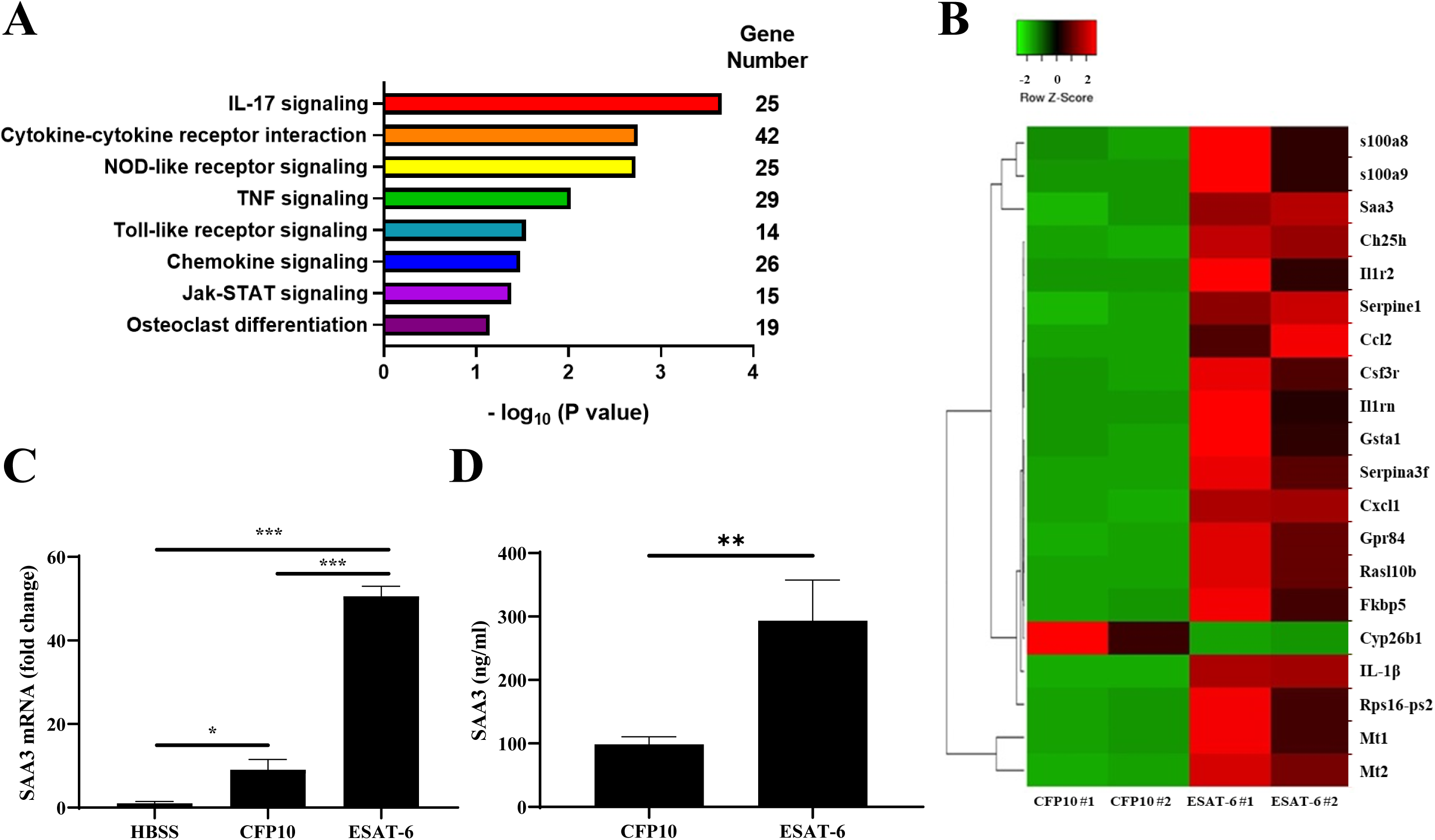
ESAT-6 stimulates the expression of SAA3 among other genes of inflammation and immune responses. C57BL/6 mice were treated with ESAT-6 or CFP10 as in Fig. 3 and the gene expression profile was determined by RNA-sequencing analysis of the total lung RNA from two mice for each treatment. The significantly upregulated pathways with gene numbers identified by gene set pathway enrichment analysis (A) and the heat map representing top 20 highly expressed genes (B) by ESAT-6 in comparison with CFP10 treatment are shown. The expression of SAA3 mRNA (C) in total RNA Of the mouse lungs and SAA3 protein (D) in mouse lung homogenates treated with ESAT-6 or CFP10 for 24 h were determined by real-time PCR and mouse SAA3 ELISA, respectively. Data expressed as Mean ± SEM. n.s., not significant; *, P < 0.05; or ***, P < 0.001.

### *Mtb* induces systemic SAA3 expression in an ESAT-6 dependent manner

To determine whether *Mtb* induces SAA3 expression in vivo in an ESAT-6 dependent manner, the expression of SAA3 mRNA and protein was determined in mouse lungs after infected with *Mtb*. Consistent with IL-1β data (Fig. 3A), H37Rv started to induce both mRNA (Fig5. A) and protein (Fig. 5B) of SAA3 at 10 DPI and induced significantly higher levels than that of H37Rv:Δ3875 at 14 DPI. Interestingly, H37Rv also induced significantly higher levels of SAA3 protein in the plasma samples of H37Rv infected mice (Fig. 5C), implying that *Mtb* induces SAA3 expression in vivo in an ESAT-6 dependent manner.

**Figure 5.**
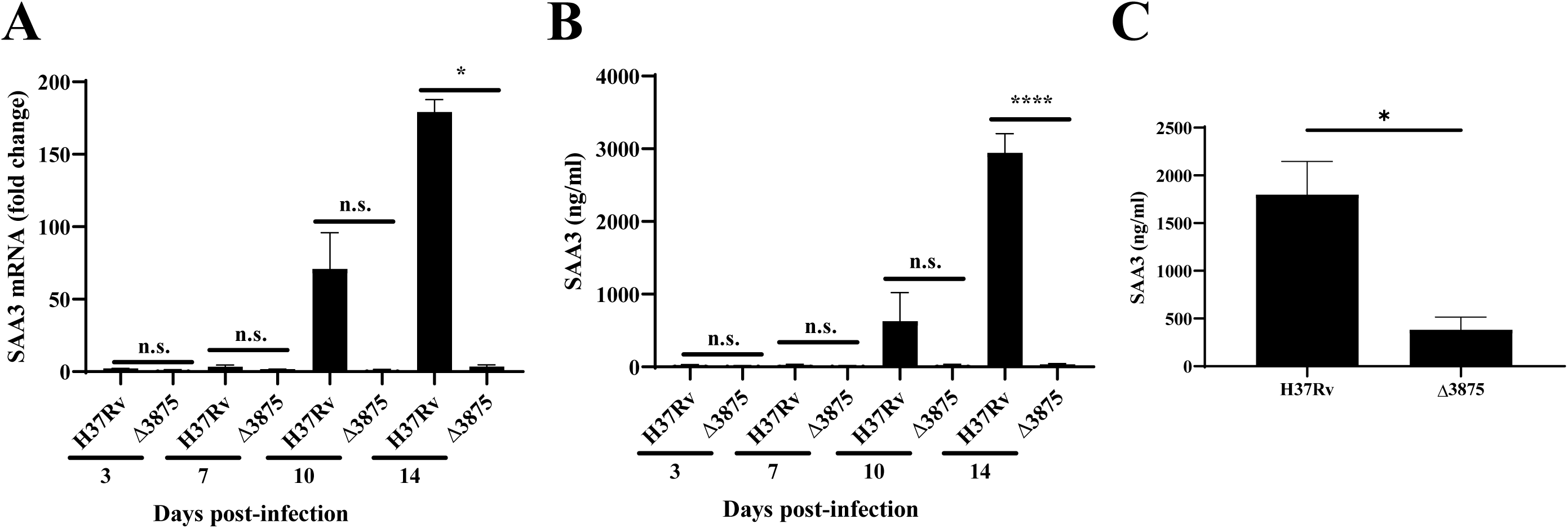
*Mtb* stimulates SAA3 expression in mouse lungs in an ESAT-6 dependent manner. C57BL/6 mice were infected with *Mtb* H37Rv or H37Rv:Δ3875 (Δ3875) intranasally as described in Fig. 3, and the expression of SAA mRNA (A) in total RNA of the mouse lung and SAA3 protein levels (B) in the lung homogenates were determined by real-time PCR and ELISA, respectively, at indicated times points post infection. The levels of SAA3 in serum samples were determined at day 14 after infection were determined by ELISA (C). Data are expressed as Means ± SEM. n.s., not significant; *, P < 0.05; **, P < 0.01 or ***, P < 0.001.

### Silencing SAA3 in macrophages leads to reduced IL-1β production by macrophages infected with *Mtb*

Our results above clearly show that *Mtb* and its major virulence factor ESAT-6 induce IL-1β production and the expression of SAA3 in mouse lung. As SAA3 was shown to induce IL-1β production by macrophages (32), we determined the role of SAA3 in *Mtb*-induced macrophage IL-1β production after silencing SAA3 expression. Silencing SAA3 in BMDMs reduced H37Rv infection induced IL-1β production significantly compared with the cells with control siRNA (Fig. 6B). The silencing efficiency was confirmed by real-time PCR for SAA3 using total RNA of siRNA transfected BMDMs in comparison with control siRNA and mock transfected BMDMs (Fig. 6A). In contrast, SAA3 silencing did not affect *Mtb*-induced IL-6 production (Fig. 6C). These data suggest that SAA3 is required for *Mtb*-induced IL-1β production by macrophages.

**Figure 6.**
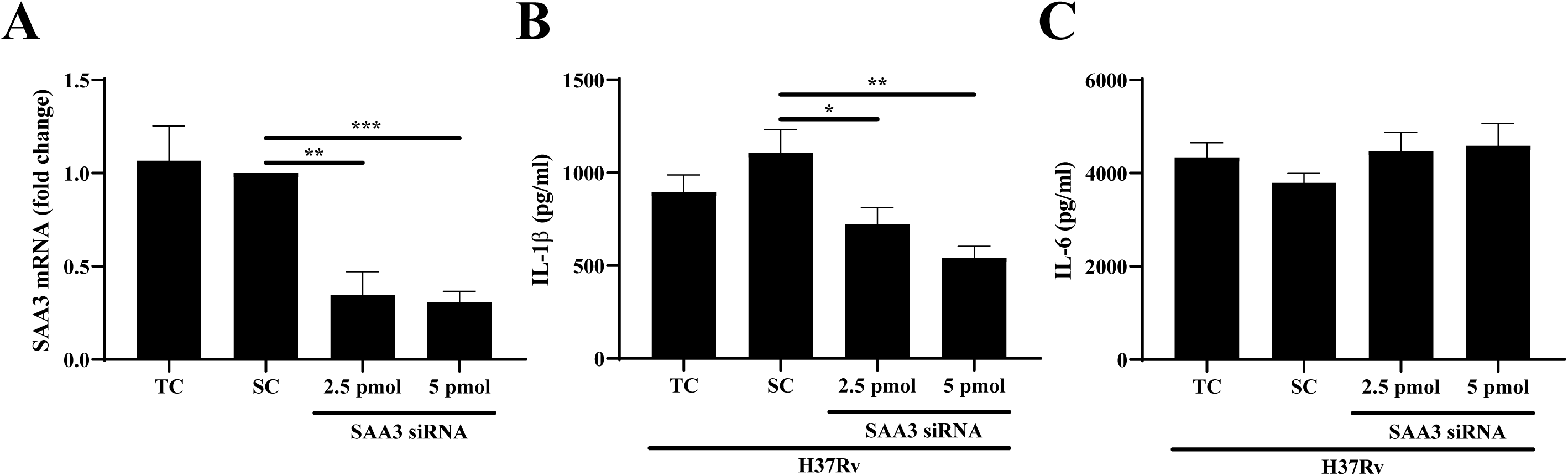
Silencing SAA3 in macrophages reduces *Mtb*-induced IL-1β production. BMDMs were transfected with the mock (TC; transfection reagent control), scrambled siRNA (SC), or two different concentrations of SAA3 siRNA for 24 h. The transfected BMDMs were infected with H37Rv at 10 MOI for 24 h. The silencing efficiency was determined by real-time PCR for SAA3 in the cells after 24 h transfection (A). The levels of IL-1β (B) and IL-6 (C) in the cell culture supernatants were determined by ELISA. Data are expressed as Means ± SEM. *, P < 0.05, **, P < 0.01 or ***, P < 0.001.

## DISCUSSION

IL-1β is essential for host protective immunity against various pathogens, including *Mtb* (3, 33, 34). However, uncontrolled production of IL-1β may cause local tissue damage due to excessive inflammation (35). A variety of human pathogens, including *Mtb* (36) stimulate the production of IL-1β by macrophages and other immune cells, such as dendritic cells and neutrophils. Previous studies have shown that *Mtb* stimulates IL-1β production by macrophages with the involvement of activated NLRP3, caspase 1 and cathepsin B in an ESAT-6 secretion system (ESX)-1 dependent manner (17, 18). In this study, we have demonstrated with data from both *in vitro* macrophage infection with *Mtb* and *in vivo* mouse intranasal treatment with ESAT-6 or infection with *Mtb* that ESAT-6 is required for *Mtb-*induced mature IL-1β production by macrophages with the involvement of K+ efflux and activated NLRP3 (17, 22). Based on this finding, we further demonstrate that ESAT-6 dependent IL-1β production by macrophages infected with *Mtb* requires SAA3, one of the major acute phase proteins associated with inflammation and immune responses. Infection of H37Rv but not its *esat-6* deletion mutant induced elevated expression of SAA3 in macrophages and mouse lungs both at mRNA and protein levels and silencing SAA3 in macrophages resulted in reduced IL-1β but not IL-6 production upon *Mtb* infection, supporting a role for SAA3 in *Mtb-*induced IL-1β production in tuberculosis infection. This is consistent with the previous reports showing the potential role of SAA3 activated NLRP3 in mature IL-1β production by macrophages in chronic allergic diseases (32) and SAA3 induces transcription and maturation of IL-1β in macrophages (14) and neutrophils (37). This study further extends the role of SAA3 in *Mtb*-induced IL-1β production, thus suggesting a potential role of SAA3 in both the immune responses against and the pathology of tuberculosis infection at least in part by regulation of IL-1β production by macrophages.

We showed that ESAT-6 deletion mutant stimulates significantly less IL-1β production by macrophages compared with that of esat-6 intact *Mtb* strains despite eliciting comparable levels of pro-IL-1β. Incubation of macrophages with ATP, NLRP3 activator, rescued the deficiency in IL-1β secretion by H37Rv:Δ3875 infected macrophages, clearly demonstrating that ESAT-6 is essential for the mature IL-1β production by *Mtb* infected macrophages, consistent with our previous report that RvD981, a MprAB two-component system deletion mutant of H37Rv, induces reduced IL-1β production by macrophages due to a deficient ESAT-6 secretion despite the expression of comparable levels of ESAT-6 to its parental strain H37Rv (22). Our in vivo studies also support a role of ESAT-6 in IL-1β production in mouse lungs upon *Mtb* infection. Thus, we speculate that ESAT-6 may function as both priming and activation signals for IL-1β production in tuberculosis infection. As a priming signal, we showed that either incubation of macrophages or intranasal treatment of mouse with ESAT-6 induced pro-IL-1β expression, consistent with our previous reports that ESAT-6 stimulates macrophage production of inflammatory cytokine and chemokine (19, 38). As an activation signal, our data with MCC950, a novel selective NLRP3 inhibitor (23, 24), showed that *Mtb* stimulates mature IL-1β production by macrophages in an NLRP3 dependent manner, consistent with the previous study [16]. We have also shown previously with human monocyte derived dendritic cells that caspase-1 is required for ESAT-6 mediated production of IL-1β (20). Therefore, the findings from this study, together with others demonstrate that ESAT-6 is required for *Mtb* infection induced activation of NLRP3 and caspase-1 for macrophage production of mature IL-1β. However, the exact details of ESAT-6 mediated activation of NLRP3 and caspase-1 remain unclear. Since localization of bacteria and their components in the cytosols of the infected cells is required for the activation of NLRP3 and caspase-1 (39), we speculate that the membrane damaging effects of ESAT-6 may contribute to the activation of NLRP3 and caspase-1 indirectly by aiding in the release of *Mtb* and its components from the phagolysosome to cytoplasm of infected macrophages (40, 41) and the candidate components with cellular NLRP3 activation remain to be identified.

Our studies with intranasal infection of mice with a high inoculum of H37Rv supported the role of ESAT-6 in *Mtb* infection induced IL-β production. Infection of mice with H37Rv induced IL-1β production in the lungs. In contrast, *esat*-6 deletion mutant did not stimulate IL-1β production, and this may be directly due to growth defect of the *Mtb* strain lacking ESAT-6. To address this directly, we administered recombinant ESAT-6 and its molecular partner CFP10 intranasally to the mice. The transcript and protein levels of IL-1β were dramatically increased in the lungs upon ESAT-6 but not CFP10 inoculation, indicating that ESAT-6 is one of the main factors of *Mtb* to induce IL-1β production in live infection.

One interesting observation from this study is that intranasal delivery of ESAT-6 into mouse followed by mouse lung gene expression profiling by RNA-sequencing demonstrated that ESAT-6 induced the expression of a large number of genes and some of which are consistent with the previously reported gene profiling studies of macrophages or the lungs of mice infected with *Mtb* (30, 31). Intranasal ESAT-6 delivery into mouse induced significant number of gene expression in mouse lungs associated with immunity and inflammation, such as IL-17 production and signaling pathways and cytokine and cytokine receptor signaling as well as the expression of wide variety of chemokine and cytokine genes, including IL-1 family cytokines. Of note, SAA3 is one of the most highly upregulated genes in mouse lungs by ESAT-6. SAA3 is a well-known DAMP to induce the expression and maturation of IL-1β by macrophages (14) and neutrophils (37) through the activation of NLRP3 inflammasome *via* P2×7 receptor and a cathepsin B-sensitive pathway (14). However, its role in *Mtb* infection in general and IL-1β production, in particular, remains unexplored. Our *in vitro* data from macrophage SAA3 silencing experiment show that SAA3 is involved in IL-1β production by macrophages infected with *Mtb*. The reduced production of IL-1β is less likely due to general defects in macrophage cytokine production because silencing of SAA3 did not affect *Mtb* induced IL-6 production by macrophages contrary to the previous reports that SAA3 is also required for IL-6 production by macrophages (42–44). This is probably due to differences in the cells with different disease settings. Based on our data and published studies of others, it is reasonable to speculate that ESAT-6 may simulate macrophage IL-1β production through elicitation of SAA3 and IL-1β, together with IL-6 may further upregulate ESAT-6 initiated production of SAA3 with a positive feedback control mechanism. This mechanism may play critical roles in tuberculosis associated chronic lung inflammation and tissue damage by helping *Mtb* establish infection in the lungs by aiding the dissemination of *Mtb* from the alveolar space into the lung parenchyma and granuloma formation (45). *Mtb* elicited SAA3 production with the involvement of ESAT-6 may also play a critical role in elicitation of Th17 immune responses in tuberculosis infection by the release of mature IL-1β by *Mtb* infected cells (46) as IL-17 plays critical roles in immune responses against tuberculosis infection (47). However, the significance of SAA3 in tuberculosis infection through regulation of IL-1β production in immune responses against and disease pathology of tuberculosis infection remains unclear and requires future studies with tuberculosis infection models of SAA knockout mouse strains.

In summary, we demonstrate that ESAT-6 plays critical roles in *Mtb* infection induced mature IL-1β production by macrophages, and SAA3 is required for *Mtb* induced IL-1β production. Therefore, targeting ESAT-6-dependent SAA3 and other members of SAA family proteins in macrophages may serve as an efficient host directed therapeutic approach for better tuberculosis control.

## Corresponding author

Buka Samten, M.D., Department of Pulmonary Immunology, University of Texas Health Science Center, 11937 US Hwy 271, Tyler, TX 75708-3154, phone (903)-877-7665, fax (903) 877-5516

## Acknowledgment

We thank Dr. Charles A. Dinarello for his thoughtful guidance and discussions.

## Brief summary

*Mycobacterium tuberculosis* stimulated the production of IL-1β by macrophages and in mouse lungs with the activation of NLRP3 and K+ efflux in an ESAT-6 dependent manner with the involvement of SAA3.

